# Deep Learning Predicts Non-Normal Peptide FAIMS Mobility Distributions Directly from Sequence

**DOI:** 10.1101/2024.09.11.612538

**Authors:** Justin McKetney, Ian J. Miller, Alexandre Hutton, Pavel Sinitcyn, Joshua J. Coon, Jesse G. Meyer

**Affiliations:** Department of Biomolecular Chemistry, University of Wisconsin-Madison, Madison WI 53706, USA; National Center for Quantitative Biology of Complex Systems, Madison, WI 53706, USA; Gladstone Data Science and Biotechnology Institute, The J. David Gladstone Institutes, San Francisco, California, USA; Quantitative Bioscience Institute, University of California, San Francisco, California, USA; Department of Cellular and Molecular Pharmacology, University of California, San Francisco, California, USA; Department of Computational Biomedicine, Cedars Sinai Medical Center, Los Angeles, CA 90048, USA; Advanced Clinical Biosystems Research Institute, Cedars Sinai Medical Center, Los Angeles, CA 90048, USA; Smidt Heart Institute, Cedars Sinai Medical Center, Los Angeles, CA 90048, USA; Morgridge Institute for Research, Madison, WI, USA; Department of Chemistry, University of Wisconsin-Madison, Madison, WI, USA

## Abstract

Peptide ion mobility adds an extra dimension of separation to mass spectrometry-based proteomics. The ability to accurately predict peptide ion mobility would be useful to expedite assay development and to discriminate true answers in database search. There are methods to accurately predict peptide ion mobility through drift tube devices, but methods to predict mobility through high-field asymmetric waveform ion mobility (FAIMS) are underexplored. Here, we successfully model peptide ions’ FAIMS mobility using a multi-label multi-output classification scheme to account for non-normal transmission distributions. We trained two models from over 100,000 human peptide precursors: a random forest and a long-term short-term memory (LSTM) neural network. Both models had different strengths, and the ensemble average of model predictions produced higher F2 score than either model alone. Finally, we explore cases where the models make mistakes and demonstrate predictive performance of F2=0.66 (AUROC=0.928) on a new test dataset of nearly 40,000 different *E. coli* peptide ions. The deep learning model is easily accessible via https://faims.xods.org.

---

Ion mobility spectrometry (IMS) has long played an important role in mass spectrometry (MS), providing an additional dimension of separation complementary to separations by liquid chromatography (LC) and mass-to-charge(Baker et al., 2007; Valentine, Counterman, & Clemmer, 1999; Valentine et al., 1998, 2006). Specifically, IMS has allowed for isolation and separation of biomolecules based on shape(Allen et al., 2017; Hale et al., 2020; Williams et al., 2020; Zhou et al., 2013), including differentiating isomers(Nagy et al., 2018; Wojcik et al., 2019), which can be a challenging task for LC-MS/MS alone. Increases in speed and efficiency of IMS, and integration of IMS with commercial MS systems has led to its widespread application in proteomics(Hebert, Prasad, et al., 2018a; Prasad et al., 2014; Purves et al., 2017; Shliaha et al., 2013; Swearingen et al., 2012; Venne et al., 2005). Among the several variants of IMS, high field asymmetric-waveform ion mobility spectrometry (FAIMS) with high transmission efficiency was commercially released in 2018 as a source-attached module, allowing for rapid separation of peptide ions on timescales compatible with LC-MS/MS experiments (Hebert, Prasad, et al., 2018b; Schweppe et al., 2019).

FAIMS relies on the differential mobility of ions in an electric field of varying strength(Cooper, 2016; Shvartsburg et al., 2006). Ions ejected from the electrospray emitter follow a roughly parabolic path through the FAIMS module before entering the mass spectrometer. Within the FAIMS module, ions pass between two electrodes where they are destabilized by an electric field with an asymmetric waveform causing their collision with one of the electrodes. An ion-specific compensating voltage (CV), set by the user, stabilizes a sub-population of ions, allowing their transmission into the instrument. In contrast with other ion mobility techniques, FAIMS therefore acts as a filter that simplifies the mixture of ions entering the MS instrument.

FAIMS filtering can simplify the gas-phase peptide ion mixture, allowing for improved detection of low abundance peptides in proteomic analysis. This increased access to low abundance peptides can improve proteomic depth, in a similar fashion to orthogonal chromatographic fractionation(Hebert, Prasad, et al., 2018a). In a process known as internal stepping, CV can be rapidly changed within a method. Such operation allows for the selection and quantification of several “gasphase fractions”, leading to an expanded number of peptide identifications without increasing sample preparation efforts. The increase in signal-to-noise and stepping speed of the FAIMS interface can also provide benefits in quantitative accuracy of targeted methods(Xia et al., 2004) and identifications in DIA strategies utilizing short gradients(Bekker-Jensen et al., 2020). In fact, FAIMS filtering has enabled fast proteome analysis without the use of LC(Meyer et al., 2020) or even multi-omics without LC(Jiang et al., 2023).

At present, the optimally transmissive CV for a given peptide must be determined empirically. Although previous work has identified some of the most important parameters affecting ion mobility, such as charge, the relationship is neither straightforward nor linear. Since the inception of IMS, researchers have investigated peptide ion behavior in hopes of developing predictive models(Valentine, Counterman, & Clemmer, 1999; Valentine, Counterman, Hoaglund-Hyzer, et al., 1999). Recent work has focused on identifying drift times(Shah et al., 2010; B. Wang et al., 2010, 2013) or collisional cross section(Meier et al., 2021). Drift time and collisional cross section are single value properties that lend themselves to regression analysis. In contrast, peptides separated by FAIMS are often found to transmit at several CVs, and CV settings tend to be designed in a discrete stepwise fashion, which makes regression less practical in this application.

Machine learning models have been applied to virtually all areas of mass spectrometry and associated technologies, seeking predictive information for a broad spectrum of molecules including proteins, lipids (Blaženović et al., 2018; Hutchins et al., 2019b, 2019a), sugars(Sabater et al., 2019), and nucleic acids (Yamankurt et al., 2019). In the proteomics field alone, substantial research has focused on the development of models predicting retention time (Afkham et al., 2017; Ma et al., 2018; Moruz et al., 2010, 2012; Pfeifer et al., 2007), fragmentation spectra (Gessulat et al., 2019; Tiwary et al., 2019), and high-signal peptides for targeted proteomic methods(Fusaro et al., 2009). Many of the resulting tools contribute to a more streamlined design of optimized methods *in silico*. These machine learning tools enable more rapid and cost-effective method development for many MS data acquisition modes (*e*.*g*., PRM, DDA, and DIA), and could be used as additional constraints for peptide identification scoring.

A recent study characterized about 60,000 peptide FAIMS transmission distributions across a range of FAIMS CVs from -25 V to -70 V(Deng et al., 2022). They computed 76 peptide properties such as quantitative structure-activity relationships from sequence features along with amino acid composition. They used these features to fit a linear model and LASSO to an output of maximum FAIMS CV (regression) and they achieved a mean absolute error (MAE) of 5.89 V. They also trained a “stacked ensemble” deep learning model to achieved at MAE of 3.81 V. These results are important, but they fail to leverage sequence context as model input, and by framing the task as regression, they do not account for what we find to be commonly non-normal transmission distributions. These non-normal distributions lead to poor predictive performance for some peptides when the problem is framed as simple regression.

Here we present a pair of machine learning algorithms, a random forest (RF)(Breiman, 2001) and a long-term short-term memory(Hochreiter & Schmidhuber, 1997) artificial neural net (NN), for *in silico* prediction of the optimal CV settings for any given peptide ion. The RF represents an approach that is both robust and easily interpretable, while the NN represents a state-of-the-art machine learning model that has shown strong performance previously in predicting peptide behavior(Meier et al., 2021). With these models and an extensive set of experimental data, we explore peptide properties that determine FAIMS transmission profiles. Although CV is a continuous parameter, regression or single label classification models do not effectively deal with the many peptides that do not show a normally distrubted pattern of transmission as a function of FAIMS CV, which often have more than one FAIMS CV with a similar maximal transmission value. Therefore, we framed this problem as multi-label classification. The ensemble based on the average of both model predictions trained on over 100,000 human peptide precursor mass-to-charge (*m/z*) peaks resulted in a 0.928 AUCROC score when predicting FAIMS CVs for a novel test set of over 40,000 *E. coli* peptide precursor *m/z* species.

## EXPERIMENTAL SECTION

### Materials and reagents

*Escherichia coli* samples of strain K12 MG1655 donated by the Cox group of the UW-Madison Biochemistry Department, were set to grow overnight at 37°C. Human lysate was derived from intact mass spec-compatible human (K562 cells) protein extract purchased from Promega (Product # V6941).

### Sample preparation for mass spectrometry

*E. coli* pellet was resuspended in 4M guanidine HCl before cells were lysed by probe sonication, boiled, and allowed to cool before being brought to 90% MeOH. Lysate was centrifuged at 15,000 g for 7 minutes and supernatant was disposed, while precipitate was resuspended in reducing and alkylating buffer (8M Urea, 40 mM TCEP, 10 mM CAA, 100 mM Tris pH 8). Lys-C was added at a ratio of 50:1 w/w protein to protease, and the sample was incubated at room temperature for 4 hours. Buffer was diluted to 25% with 100mM Tris pH 8 and trypsin was added at a ratio of 50:1 before the sample was digested at ambient temperature overnight. Human extract was prepared in an identical manner starting with addition of methanol. Digested peptides were desalted using Strata-X Polymeric Reverse Phase column (Phenomenex).

### nLC-MS/MS data collection

Online reverse-phase columns were prepared in-house using a high-pressure packing apparatus(Shishkova et al., 2018). In brief, 1.7 µm Bridged Ethylene Hybrid C18 particles were packed at 30,000 psi into a New Objective PicoTipTM emitter (Stock# PF360-75-10-N-5) with an inner diameter of 75 µm and an outer diameter of 360 µm. During separations, the column was heated to a temperature of 50 °C inside a heater (developed in-house) and interfaced with the mass spectrometer via an embedded emitter.

A Dionex UltiMate 3000 nanoflow UHPLC was used for online chromatography with mobile phase buffer A consisting of 0.2% formic acid in water and mobile phase buffer B consisting of 0.2% formic acid in 70% acetonitrile. Mobile phase B was increased to 9% in the first 6.5 min then increased to 43% B at 41 min. The method finished with a wash stage of 100% B from 44-48 minutes and an equilibration step of 0% B from 50-60 minutes, for a total method length of one hour. Flow rate was 335 nanoliters per minute throughout.

Eluting peptides were ionized by electrospray ionization and passed through the FAIMS Pro module (Thermo Scientific) using a single compensating voltage for the duration of each method. Ions were analyzed on a Thermo Orbitrap Fusion Lumos with survey scans taken from 300 to 1500 m/z at 240,000 resolution while using Advanced Precursor Determination(Hebert, Thöing, et al., 2018) and an AGC target of 1e6 with a maximum injection time of 50 ms. Precursor isolation used a window of 0.7 Th with 15 ppm mass tolerance and a dynamic exclusion time of 20 seconds. Selected precursors were fragmented using HCD with a normalized collision energy of 25%. The MS2 AGC target was set at 3e4 with a maximum injection time of 14 ms and scans taken using the turbo setting. Analysis was performed in technical duplicate.

### High Resolution FAIMS data with 1-volt steps

Additional samples from *E. coli*, yeast, C elegans, and Human were prepared similar to as described above. The data was also collected similar to as described above with the key change that “High Resolution” FAIMS mode was used, which sets the inner electrode temperature to 70°C instead of 100 °C, and steps of 1 volt were used for CV from -15 to -90. The human data was analyzed by MaxQuant, and the peptide transmission distribution was produced from the evidence.txt file.

### Peptide Identification and Quantification

Raw files were converted to .mzML format using Proteowizard(Kessner et al., 2008) (version 3.0.19039). Peptides were identified by database search against the human proteome including isoforms (downloaded 2019-12-19) or the *E. coli* proteome (downloaded 2020-05-01) using MS-Fragger(Kong et al., 2017) (version 2.2) through the FragPipe GUI (version 12.1). The default search parameters were used except for the precursor tolerance, which was set to +/-5 ppm and the fragment tolerance was set to 0.5 Daltons. Database search output files in pep.xml format were combined into one file using iProphet(Shteynberg et al., 2011) within philosopher(da Veiga Leprevost et al., 2020) (version 2.0). Peptide identifications were imported into Skyline(MacLean et al., 2010) for quantification using an iProphet score cutoff of 0.99. In an attempt to determine the true distribution of peptide precursor transmission through the FAIMS device independently of the stochastic DDA identification process, precursor (MS1) signals were extracted from all files corresponding to FAIMS CV values from -20 V to - 100V. MS1 signal was extracted within 8 ppm of the predicted precursor mass and 1 minute (*E. coli* data) or 0.5 minutes (human data) of the MS/MS identifications. Precursor signals for decoy peptides were also extracted to allow mProphet determination of statistical significance of peak peaking(Reiter et al., 2011). Only precursor ion peaks with q-value of < 0.01 were integrated, and a custom Skyline report was output containing the q-values, precursor ion dot product with the theoretical isotope envelope (idotp), and peak area for each FAIMS CV. The skyline report was further processed in R to determine the FAIMS CV transmission profile for each peptide precursor ion.

### Determination of peptide CV transmission profiles

Skyline reports were read into R (version 3.6.3) and decoy peptides along with peptides matching reversed proteins were removed. Each peptide’s intensities were scaled from 0-1, with 1 representing the CV setting with maximum intensity and therefore, transmission (CVmax).

Labels were assigned based on a peptide’s CVmax. A CV bin was assigned a positive label if it was greater than both 0.5 and the average proportional intensity of the second-best CV setting for peptides sharing that CVmax (**Supplemental Figure 1**. For example, if the max CV setting for peptide X was - 45 V, because the second-best setting for these peptides is -50 V with an average of 0.52, an individual setting would be required to be >0.52 to receive an additional positive label. This labelling scheme allowed for more conservative label prediction with a focus on the top 1-3 labels, additional discriminatory ability regarding co-occurring labels, and a compensating effect for peptide groups with broader distributions. Finally, peptides with non-consecutive labels were removed from the dataset, as they were considered reflective of potentially spurious identifications or stochastic events that may undermine the predictive performance of the model. This group included <5% of all peptides. The R script (Data/human/Preprocessing_FeatureMaxCVIsolation.R) for class labeling is available on GitHub (https://github.com/Justin92/FAIMS_MachineLearning).

### Assignment of True Labels and Accounting for Label Imbalances

Initially, true positive labels were assigned to all settings with proportional intensity >0.5. This labelling scheme led to common co-labeling of peptides transmitted at higher magnitude voltages (-45 to -95) due to their broader transmission distribution (**Supplemental Figure 1**), including ions with seven or more labels. In hopes of increasing discriminatory ability, labels were assigned based on a peptide’s highest intensity CV (CV_max_). A CV bin was assigned an additional positive label if it was greater than both 0.5 and the average proportional intensity of the second-best setting for peptides sharing that CV_max_. This labelling scheme allowed for more conservative label prediction, with a focus on the top 1-3 labels discussed above, and a compensating effect for peptide groups with broader distributions. This labelling scheme also reduced the number of ions with 6 or more labels by >30%.

Label imbalances make classification tasks difficult(He & Ma, 2013), which can be compounded in multilabel schemes(Charte, Rivera, del Jesus, et al., 2015; Charte, Rivera, Del Jesus, et al., 2015; Tahir et al., 2012) due to the dominant proportion of negative labels. In our full human peptide dataset (128,000 peptide ions) our rarest label (CV=-20) includes 727 peptides, composing <1% of the peptide ion population. These imbalances lead to substantial underprediction from our random forest model across all labels with >40% peptides receiving no predicted labels (**Supplemental Figure 2**). A common solution in the field involves oversampling positive labels through resampling or addition of synthetic training data(Han et al., 2005; He et al., 2008; Zheng et al., 2016). To increase the number of predicted labels by the random forest, we allowed the model to see more of each label class during training by lowering the proportional intensity threshold used to assign additional labels in the training data without changing the test set label threshold. This increased training label number, leading to an increase in predicted labels and predictions that more closely matched our true labels to improve recall and F2 score (**Supplemental Figure 2**) at the minor expense of thresholded accuracy (**Supplemental Table 1**).

Three labelled peptide groups were utilized in our analysis. Two were derived from the human peptide dataset, a 70% training set and a 30% test set to assess generalization performance. The third group included all non-overlapping peptides (99.7%) from the *E. coli* dataset for external evaluation of the models’ performance. The human dataset was split in an iterative fashion that preserved label distributions between the full and split datasets (**Supplemental Figure 3**) using the IterativeStratification function in scikit-multilearn(Szymański & Kajdanowicz, 2017) (version 0.2.0). The human 70% training set was used for optimizing the model hyperparameters. After optimizing the models with the 70% human training data, final models were trained using the full human peptide dataset, and the full model was used to make predictions against the true test set of non-overlapping *E. coli* peptides.

### Machine Learning - Random Forest

The random forest (RF) model received as inputs: peptide length, charge state, mass, isoelectric point (pI), and one-hot encoded sequence. Length and charge were both derived from the original peptide search results and labelling process. Mass and pI were generated using the pyteomics(Goloborodko et al., 2013; Levitsky et al., 2019) (version 4.3.3) package in Python based on the unmodified sequence of the peptide. The sequence was one-hot encoded using Keras (version 2.2.4) with 22-member alphabet that included the 20 standard proteinogenic amino acids as well as the common modifications N-terminal acetylation and oxidation of methionine. The multilabel random forest classifier was developed based on the use of the On-eVsRestClassifier wrapper function in tandem with the Ran-domForestClassifier function in scikit-learn (Pedregosa et al., 2011) (version 0.23.0) in Python. Although all input features improved performance for some labels, charge and one-hot sequence were crucial to performance of the model (**Supplemental Figure 4A**).

Three different strategies were used to optimize the hyperparameters for the random forest: a parallel grid search, a random search using the RandomizedSearchCV function from scikit learn, and a Bayesian search using the hyperopt(Bergstra et al., 2013) package (version 0.2.4) in Python. Hyperparameter optimization focused on five parameters: (1) number of trees in the forest(“n_estimators”), (2) the maximum depth of the trees(“max_depth”), (3) the number of features to use at each split(“max_features”), (4) the minimum number of samples required at each leaf node (“min_samples_leaf”), and (5) the minimum number of samples required to split a node (“min_samples_split”). The grid search tested 2191 total hyperparameter combinations (**Supplemental Table 2**) largely in parallel using the compute resources and assistance of the UW-Madison Center for High Throughput Computing (CHTC) in the Department of Computer Sciences. Due to computational limitations and the sequential nature of the random and Bayesian search, each was run for 50 iterations, with weighted ROC AUC used as the scoring metric. Despite the abundance of local maxima and minima even when varying only two hyperparameters (**Supplemental Figure 4B**) Bayesian and random search both efficiently navigated the search space. The Bayesian optimization used knowledge of prior hyperparameter sets to more efficiently utilize computing resources with a greater number of models tested in productive feature space (**Supplemental Figure 4C**). In the interest of balancing recall and precision, a hybrid set of hyperparameters was used, combining aspects of the Bayesian and grid optimization. The final set of hyperparameters utilize: (1) 450 trees, (2) a maximum depth of 30, (3) 0.3807 of the features at each split, (4) a minimum of 6 samples per leaf node, (5) a minimum of 9 samples to split a node. The given hyperparameters for any machine learning problem can be infinitely iterated, but we have identified here a strong hyperparameter set that significantly increased performance as compared to the default hyperparameters. The Python code for all three hyperparameter optimizations is available on GitHub. (https://github.com/Justin92/FAIMS_MachineLearning).

To mitigate the label imbalance between the most common CV labels and the rarest, training data for the random forest was labelled using lowered thresholds for positive labels (**Supplemental Figure 2**). These thresholds are described in **Supplemental Table 3** and were applied to the 70% human training data when testing against the 30% test set (**Table 1**). These thresholds were also applied to the full human data when training the random forest for validation against the *E. coli* peptide dataset (**Table 2**).

**Table 1.** Performance metrics from human validation dataset. Binary accuracy, precision, recall, f2 and ROC AUC are all calculated when training each model and an averaged ensemble using 70% of the human data and testing against the remaining 30%. At least one match describes the proportion of peptides that share at least one positive label between experimental labels and predicted labels.

| Metric | Random forest | Neural network | Ensemble |
| --- | --- | --- | --- |
| Binary accuracy | 0.8133 | 0.9073 | 0.8970 |
| precision | 0.3900 | 0.6610 | 0.5774 |
| recall | 0.8271 | 0.5581 | 0.7136 |
| F<sub>2</sub> | 0.6756 | 0.5760 | 0.6814 |
| ROC AUC | 0.8977 | 0.9414 | 0.9344 |
| At least one match | 0.9018 | 0.7535 | 0.8403 |

**Table 2.** Performance metrics from the *E. coli* test data. Binary accuracy, precision, recall, f2 and ROC AUC are all calculated when training each model and an averaged ensemble using 100% of the human data and testing against the full *E*. coli precursor dataset. At least one match describes the proportion of peptides that share at least one positive label between experimental labels and predicted labels.

| Metric | Random forest | Neural network | Ensemble |
| --- | --- | --- | --- |
| Binary accuracy | 0.8023 | 0.9027 | 0.8950 |
| precision | 0.3789 | 0.6503 | 0.5830 |
| recall | 0.8084 | 0.5487 | 0.6837 |
| F<sub>2</sub> | 0.6590 | 0.5664 | 0.6609 |
| ROC AUC | 0.8834 | 0.9368 | 0.9280 |
| At least one match | 0.885 | 0.7447 | 0.8195 |

### Machine Learning - Neural Network

The LSTM NN was built with the keras, which is a high-level interface to tensorflow (version 2.1). The exact anaconda environment can be created using the file on the github repository at neural_net/tensorflow21.yml. Precursor charge was appended to the front of the peptide sequence, and the string of charge and sequence was converted to a string of integers for input to the neural network. Peptides with length less than 50 were padded with an integer encoding “end” to ensure all inputs were the same length. Neural network output was a string of 16 probabilities corresponding to each of the possible CV values from - 20 volts to -95 volts in steps of -5 V. A positive class was assigned for any prediction 0.5.

The general neural network structure used four layers with two dropout steps and one batch normalization: (1) an embedding layer that converted the length 51 integers into vectors of real-value numbers, (2) the LSTM layer, (3) dropout, (4) dense with ReLU activation, (5) dropout, (6) batch normalization, (7) dense output with sigmoid activation. Binary crossentropy loss was used with the Adam optimizer. Keras-tuner was used to optimize the LSTM hyperparameters targeting high ROC AUC on the validation set with the hyperband algorithm. The hyper-parameter optimization was done using 5-fold cross validation within the 70% human training data slice. Possible hyperparameters were: (1) number of outputs from the embedding layer, from 32 to 96 in steps of 32, (2) number of outputs from the LSTM layer from 32 to 96 in steps 32, (3) dropout proportion from 0 to 0.5, (4) number of outputs from dense layer 1 from 32 to 96 in steps of 32, (5) dropout proportion from 0 to 0.5, (6) learning rate from 1e-4 to 1e-2 with log sampling. The optimal parameters were: (1) embedding layer output of 96, (2) LSTM output of 96, (3) dropout #1 of 0.32255, (4) dense output of 64, (5) dropout #2 of 0.02152, (6) learning rate of 0.001758. The best hyperparameters were then used to find the best number of epochs with early stopping monitoring the validation set ROC AUC in 5-fold cross validation, and the 52 epochs was best. A model was then trained using all the 70% human training data, and the performance was evaluated using the held-out 30% human test set. A second model was then trained using all 100% of the human peptides, and this model was evaluated with the second external test set of ∼40,000 peptides from *E. coli*.

### Data and Code Availability

Raw mass spectrometry data and MS-Fragger search outputs are available from the Mas-sIVE(M. Wang et al., 2018) repository (massive.ucsd.edu project MSV000085707 https://doi.org/doi:10.25345/C5X756) or Pride (PXD021174). The high resolution FAIMS data is found at Pride (Project accession: PXD055252, Username, Password: XYV3wB5wa5p8) Skyline documents containing quantitative data for each peptide are available on Panorama (Sharma et al., 2014) (https://panoramaweb.org/AOueYU.url). The skyline reports, processed data, and machine learning code is available on GitHub (https://github.com/Justin92/FAIMS_MachineLearning).

We setup a website to offer an accessible interface for the neural network model (https://faims.xods.org). This model is not exactly the same as the results presented in the paper because of package incompatibilities. We refit the model with a newer version of tensorflow that is compatible with Flask, and the new model produced highly similar metrics on the *E coli* test data. The server is running on an Amazon EC2 instance using the Flask framework. From the main page, users can submit a list of peptides where it will be automatically converted into a suitable format for the model. The model is executed on each peptide, and the predictions are returned to the user as a list of predicted CV labels, a CSV file for the value of predictions, and a bar plot showing the distribution of predictions across peptides if the peptide list is sufficiently short. No login is required to use the model; results are associated with the user’s browser session. The code underlying the site is available from https://github.com/xomicsdatascience/faims.

## RESULTS

To determine optimal CV values for thousands of peptide precursor ions we first digested complex protein mixtures of both human and *E. coli* cellular lysates. These tryptic peptide mixtures were separately analyzed by capillary nano-liquid chromatography coupled with a FAIMS-enabled tandem mass spectrometer (nHPLC-FAIMS-MS/MS). To obtain high resolution CV profiles, we acquired single shot nHPLC-FAIMS-MS/MS experiments at defined CV values ranging from -20 to -100V with 5V increments (**Figure 1A**). In total we collected 20 single shot datapoints for each cell lysate. These data files were searched using MS-Fragger and filtered to an FDR of 1%. Identified peptide ions were then quantified using Skyline with an iProphet score cutoff of 0.99. Missing values were accounted for by matching MS1 features within 8 ppm and a 1-minute retention time window. Those matches were then filtered using a q-value cutoff of 0.01 based on mProphet feature analysis. After processing, a total of 128,402 and 42,719 unique peptide ion *m/z* peaks from human and *E. coli* tryptic digests, respectively. In total these spectra were mapped to 83,584 and 29,996 unique peptides sequences detected from human and *E. coli*, respectively. Note most peptides had precursor m/z values occurring in multiple charge states with the distribution being – 56.7% (+2), 35.1% (+3), 7.1%(+4) and 1.1%(+5) (**Figure 1B**).

**Figure 1.**
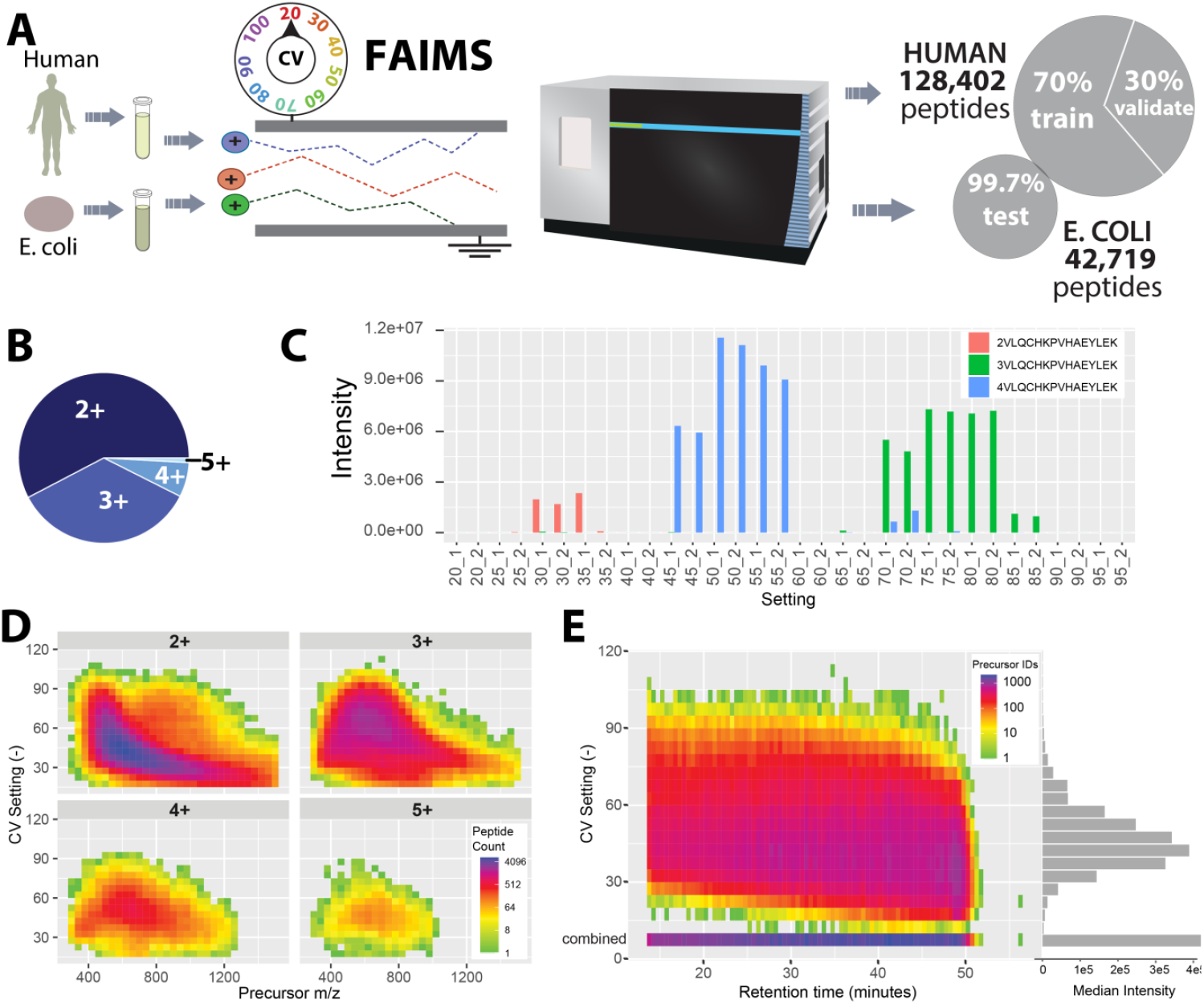
Experimental and data overview. (A) Experimental Design. Human and *E. coli* protein lysate were denatured and digested with trypsin and lys-C before being analyzed using a series of data-dependent single-CV one-hour analyses. More than 120,000 and 40,000 peptide precursor ions were quantified for Human and *E. coli* samples, respectively. (B) Pie chart depicting the proportion of identified peptides from each charge state for all human peptides, a majority of which are 2+. (C) Intensity distribution for three charge states of example peptide VLQCHKPVHAEYLEK across experimental CV range with replicates shown. Different charge states exhibit different CVmax as well as different transmission distributions. (D) Density plot of number of human peptides observed at each charge state and compensating voltage across the mass-to-charge range. Observe distinct distributions by charge state. (E) Density plot depicting number of human peptides observed at each compensating voltage for the duration of the method, sorted into 0.5 minute RT bins. Horizontal bar plot depicts the median peptide intensity at each compensating voltage. Combined indicates number of unique human peptides identified across all FAIMS CV experiments at each point in the gradient, with the horizontal bar representing the median intensity of all identified human peptides.

This high-resolution FAIMS CV data from 5V steps showed that although transmission of individual peptides can be quite variable, in aggregate, peptides exhibit substantial transmission (>50% of max) across 1-3 of the 5V bins tested here. When grouping peptides by their most transmissive CV (CV_max_), groups exhibit different average transmission distributions. Peptide ions with maximum transmission at lower magnitude CVs tend to have narrower shapes, while those that favor higher magnitude voltages exhibit wider distributions. Although in aggregate the peptides appear to have a normal distribution (**Supplemental Figure 5**), individual cases are often much more irregular in transmission distribution shape (**Figure 1C**). This variability and frequent non-normality in distribution shapes motivated our multilabel approach.

To further validate this observation of non-normality we performed two additional tests (**Supplemental Figure 6**). First, we looked at peptide transmission distributions over FAIMS CV space from Deng *et al*. We again found that these distributions were commonly non-normally distributed. Second, we collected additional data with “high resolution” FAIMS mode at a step size of 1 V instead of 5 V. This high resolution FAIMS setting will compress and shift distributions making them not directly comparable with standard resolution, but this should give the best picture of transmission distribution shapes. Examining the same peptide as **Figure 1C**, we found that in fact the +3 and +4 charge states have multiple maxima and non-normal transmission distributions (**Supplemental Figure 6C**). This may result from multiple gas phase ion structures that result in slightly different mobilities.

Further examining the broad trends in peptide transmission, we observed charge state to be the most important factor in deciding the distribution of peptides across the CV space. This relationship between peptide or protein charge state and differential ion mobility has been widely reported in many types of ion mobility(Meier et al., 2021; Shah et al., 2010; B. Wang et al., 2010, 2013) including FAIMS(Hebert, Prasad, et al., 2018a). Although there appears to be a broadly inverse relationship between peptide charge and CV, it is not linear, and higher charge states exhibit narrower CV distributions. For example, 90% of charge state 5 peptide ions were observed between CV -45 V and -55 V, with -50 being the most populated bin, whereas only 23% of charge state 2 peptides are observed in this range (**Figure 1D**).

FAIMS can enhance detection of low abundance peptides by filtering out high abundance components of complex peptide mixtures. At more extreme CV settings, we observed a greater simplification effect, as indicated by reduced peptide observations. When overlaying all peptides from across the full CV range we identify >2000 peptides per minute for much of the LC gradient (**Figure 1E**, “combined”) indicating the increased depth provided by FAIMS. Further supporting the theory that greater filtering leads to increased detection of low abundance peptides, we observed decreased median intensities at the edges of our measured CV range (**Figure 1E**). Median intensity largely mirrored peptide observations across CV space with a maximum at -45 V and minimum at -120 V, the most and least populated CV bins, respectively.

Based on the data shape we aimed to develop a model that would take peptides as input and output transmissive CVs. Although compensating voltage is a continuous variable, its discrete nature here led us to a classification model where peptides would be given a label for each compensating voltage included in our experimental dataset (5V bins from -20 to - 95). Experimental data was collected to -110 but very few peptides were identified between -95 and –120. A positive label (“1”) would indicate substantial transmission at that CV setting, while a negative label(“0”) would indicate poor or no transmission. We selected a multioutput multilabel classification framing with the objective of identifying the 1-3 most transmissive compensating voltage settings. Two technical challenges arose from this multi-label strategy and objective: (1) converting continuous intensity into a binary label indicating substantive transmission at an individual setting and (2) Compensating for large label imbalances commonly found in classification problems(Fernández et al., 2018). This first challenge was overcome by adopting a labelling scheme that identified true positive labels based on intensity relative to the peptide max and the peptide transmission distributions (**Supplemental Figure 1, Materials and Methods**). The second challenge, label imbalances, are a common issue in classification tasks(He & Ma, 2013), and can be compounded in multilabel schemes(Charte, Rivera, del Jesus, et al., 2015; Tahir et al., 2012) We addressed this issue by increasing the leniency for rare labels in our training data, a process described in detail in the methods (Material and Methods, **Supplemental Figure 2, Supplemental Table 3**).

The human peptide data was split into 70% training set and a 30% testing set based on multi-class membership with scikit-multilearn(Szymański & Kajdanowicz, 2017)(Materials and Methods) (**Figure 1A**), which preserved the proportional distribution of labels (**Supplemental figure 3**). Both models were effective at predicting optimal CV settings (**Table 1**) with the RF and NN producing binary accuracies of 0.81 and 0.90, respectively, compared to 0.49 for a uniformly random dummy classifier. The NN showed greater precision (NN = 0.66 vs. RF = 0.44) and reduced false positives, resulting in a greater ROC AUC score (0.94 vs. 0.86). The RF model favored selection of a greater number of relevant elements at the expense of false positives leading to increased recall (RF = 0.83 vs. NN = 0.56) and F2 score (0.70 vs. 0.58).

One utility of predicting FAIMS CV is for targeted method design. Given the continuous nature of compensating voltage, perfect matches between the profiles of predicted and experimental settings are uncommon. However, even a reduction in the possible CV range from 100 V (twenty possible) to 20V (four possible) would be useful in experimental design. With this utility in mind, we quantified the proportion of peptides in which at least one label was predicted correctly. We found that 75% and 90% of peptides had at least one label in common between the predicted and experimental labels in the neural net and random forest, respectively.

*E. coli* labels were assigned in the same manner as the human test data labels (Materials and Methods). When both models were trained using the full human peptide data set and validated against an entirely naïve group of *E. coli* peptide ions, overall prediction metrics were quite similar, with only minor declines in performance (**Table 2**). This similar performance on peptides that were never seen during training indicated a lack of overfitting and good generalization performance. This performance also suggested the recognition of underlying peptide properties by both algorithms and the application of these properties to transmission predictions.

Receiver operator characteristic curves (ROC-AUC) for *E. coli* label predictions help to visualize the per-CV performance of the models (**Figure 2**). Both models exhibit variable performance across the different CV labels. Interestingly, the algorithms exhibit greatest area-under-the-curve for labels with fewer observations, approaching unity in the ROC curve for -25 and -20 CV. The models struggle most within the middle voltage range (-45 to -55), despite a wealth of examples in the training data. This may be caused by the peptides in this range commonly having >3 labels, leading to difficulty in identifying discriminatory properties.

**Figure 2.**
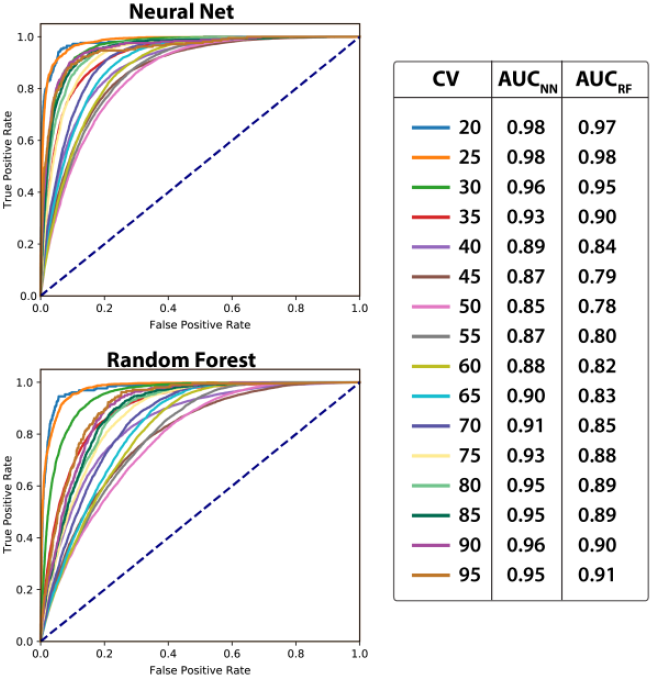
Receiver Operator Characteristic Curve for E. coli Predictions. Receiver operator characteristic (ROC) curve for each compensating voltage label in each of the two models, the neural net and the random forest when predicting labels for E. coli peptides. Observe slightly better area-under-the-curve (AUC) from neural net across all CV labels. Both models exhibit best performance on voltages closest to zero.

To better understand the prediction qualities of each model, we picked illustrative peptide ion examples from the *E. coli* dataset with true labels spanning a wide range of CV space. This analysis revealed that the distribution of CV prediction probabilities mirrored the true intensity distributions (**Figure 3**). These individual peptides examples highlight the differences in metrics - described in aggregate above - between the two models. The neural net provides a more selective set of labels, while the random forest generates wider distributions with higher probabilities, resulting in more false negatives and false positives for the NN and RF, respectively. Despite these differences in label predictions, both models reflect the asymmetrical transmission distributions that broaden as the voltage increases in magnitude. These examples illustrate the capacity for machine learning to parameterize the chemo-physical properties of peptides to predict their ion mobility in CV space.

**Figure 3.**
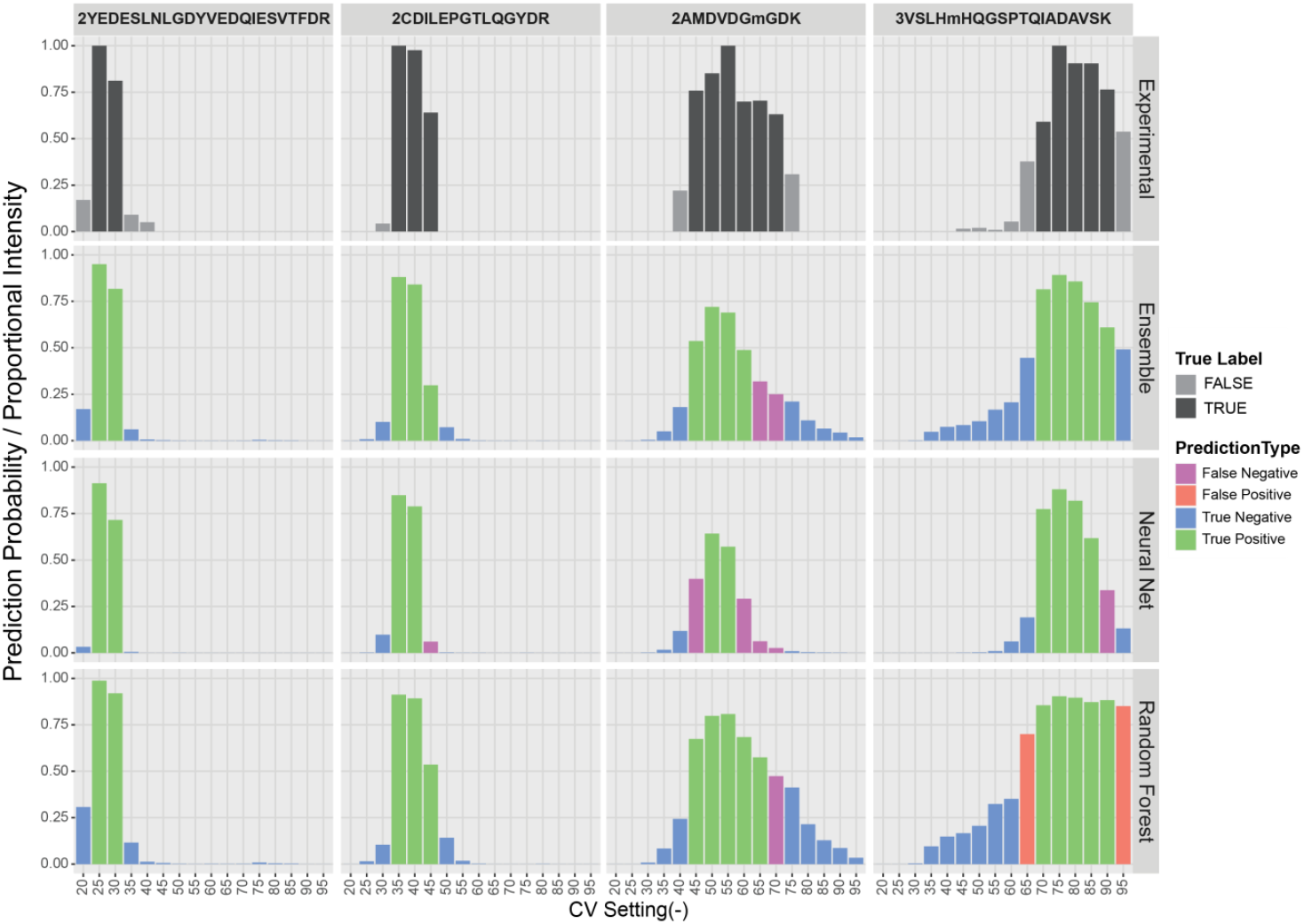
Example Peptide Prediction Probabilities. Prediction probabilities and proportional intensities for the tested CV settings for each of four example peptide ions: YEDESLNLGDYVEDQIESVTFDR +2, CDILEPGTLQGYDR +2, AMDVDGM(Ox)GDK +2, and VSLHM(Ox)HQGSPTQIADAVSK +3. For experimental data, dark grey indicates positive labels and light grey indicates negative labels. For predictions, green indicates true positives blue indicates true negatives, red indicates false positives and purple indicates false negatives.

Based on this observation of complementary predictions, we investigated if an ensemble of the model’s predictions would provide improved performance. The class predictions for each model were averaged, and the same metrics were re-computed using the average probabilities (**Table 1, 2**). Nearly all the metrics fell between the original values of the two separate models. However, F2 score, which summarizes both precision and recall as the harmonic mean, was higher for the ensemble than for either model alone, suggesting a benefit in combining the two separate prediction strategies.

To better understand what causes the models to be incorrect, an incorrect prediction was traced back to the raw data (**Supplemental Figure 7**). This peptide was predicted to transmit through FAIMS at CVs -55, -60, -65, and -70, but was assigned true labels of -50 and -55. Inspection of the precursor ion areas across the individual CV runs appears to show high signal in CV settings of -50 and -55 as the peptide was labeled, but there was an apparent low level of precursor signal in some of the predicted channels with higher precursor shape matches to the theoretical envelope pattern (idotp, as generated by Skyline). A view of the extracted ion chromatogram (XIC) from the CV of -55 shows one prominent peak with a much smaller peak in the future by 0.3 minutes. Further inspection of the precursor isotope pattern of the more prominent peak from the XIC in the “true” label CV -55 analyses showed that the incorrect peak was chosen; clearly the M+1, M+2, and M+3 peaks of a different peptide were integrated. After correcting the integration to the second, smaller peak, the distribution of integrated peaks better matched the values predicted by the neural network. Therefore, in this case, the model predictions were more accurate than the “true” labels. This highlights the importance of generating ground truth data for model training, as well as the difficulty of data labeling for this task of peptide ion mobility through FAIMS.

We demonstrate here an effective ensemble model for predicting transmissive CVs for peptides based on sequence. FAIMS has been shown to increase protein identifications in global abundance analyses and increase accuracy and precision in targeted experiments, such as PRM. The algorithm described here can be used to design these type FAIMS-MS/MS experiments optimized for specific proteomes, tissues, or target transitions without the need preliminary experiments, requiring additional material and time.

## DISCUSSION AND CONCLUSIONS

FAIMS offers an opportunity to increase proteomic depth and sensitivity without increasing sample preparation time. However, the promise of improved targeted sensitivity, further developments in targeted FAIMS methods, and new ideas for discrimination of true and decoy matches is currently slowed by necessary method optimization experiments. We present here a pair of machine learning models that when applied individually or in tandem allow the circumvention of the empirical FAIMS method development process by predicting optimal compensating voltages for peptides of interest. The models display excellent predictive performance when validated against a naïve collection of *E. coli* peptides exhibiting 0.88 and 0.93 AUC-ROC for the random forest and neural net, respectively. The models also show the capacity to approximate the transmission distributions of individual peptides across voltage range used here.

The multilabel classification strategy utilized here represents a substantial divergence from the regression-based algorithms commonly used for ion mobility behavior prediction. This strategy capitalizes on the distinct transmission distribution of individual peptides and interprets and predicts a two-dimensional shape. Although we believe multi-label classification was the optimal approach to frame the problem, this strategy introduced several technical challenges, including variable CV transmission profiles and label imbalances. We adapted our labelling scheme to provide the most useful prediction information, which we decided was a small set of optimal CV settings rather than a large range.

The models performed well when tested against model-naïve set of *E. coli* peptide ions, with the strongest performance observed for CV bins with the fewest training examples. This performance suggests generalizability of these models to tryptic peptides regardless of sample source without a need for additional data collection. Interestingly, the models performed worst when making peptide transmission predictions for the most populous voltage range of the training data. This decreased performance indicates that although we have the capacity to survey a greater number of peptide ions, it may not be beneficial to predictive accuracy.

When combining peptide identifications across our entire voltage range, we identified more than 120,000 human peptide precursor ions using a one-hour LC gradient method. These peptides represent the peptide space accessible to a FAIMS-integrated mass spectrometry method. Our CV prediction methods allow for access to this vastly expanded peptide space for any organism or any tissue. These models add to the ever-expanding toolkit for developing LC-MS methods *in silico*, which includes predictions for spectral fragmentation and retention time. When used in tandem, these tools allow for greater depth in untargeted experiments (e.g. DIA) and increased sensitivity and quantitative accuracy for targeted experiments (e.g. PRM). This could be especially useful for challenging tissues or cell types, such as plasma or cerebrospinal fluid.

As an example of how this prediction tool could be utilized, we entered the coronavirus peptide +2QQTVTLLPAADLDDFSK around which Renuse and colleagues recently developed a targeted parallel reaction monitoring method using FAIMS for rapid diagnosis of COVID(Renuse et al., 2020).Our ensemble model would have classified voltages -30 and -35 as transmissive settings, overlapping with their chosen CV of -30, without any need for optimization experiments. Increasing development speed in this way can be crucial in diagnostic development especially in the case of infectious diseases.

Many parameters, such as those associated with chromatographic gradients, electrospray ionization, mass spectrometry instruments and data searching, can affect identification and quantification of peptide ions in data-dependent experiments. Despite these parameter differences, when comparing overlapping peptides from our human dataset to those recently collected by Bekker-Jensen and colleagues using FAIMS and substantially shorter gradients(Bekker-Jensen et al., 2020), we observe similar transmission patterns (**Supplemental Figure 8**).

Although our analysis here focuses on largely unmodified tryptic peptides, based on our success, we expect ions from modified peptides or metabolites will be amenable to prediction of FAIMS transmission. Our human peptide dataset included N-terminal acetylation and oxidation of methionine, increasing our peptide sequence alphabet to 22. The available residue alphabet could easily be expanded to include other commonly modified residues allowing for predictions of a wider peptide space. Given the baseline of these effective models, retraining for modified peptides and non-tryptic peptides should be straightforward given a set of at least 10,000 additional training examples. We expect that our success in modeling peptide transmission through FAIMS could potentially be replicated for lipids or polar metabolite transmission profiles. Our machine learning modeling results more generally will also inform future machine learning studies that use peptides as input. Altogether, we expect these tools to be widely adopted in studies that utilize FAIMS. The deep learning model is accessible through a simple web interface at https://faims.xods.org

## ASSOCIATED CONTENT

### Supporting Information

Table S1 (Performance metrics for 5-fold cross validation for true and tuned labels); Table S1 (Random Forest Grid Search Hypar-parameters); Table S3 (Updated label thresholds for RF positive class training); Figure S1 (Average proportional intensity grouped by CV_max_); Figure S2 (Optimization of features and hyperparameters); Figure S3 (Mapping label combination); Figure S4 (Alternate training label threshold for random forest); Figure S5 (Examples of incorrect automatic peak picking effect on predictions).

## AUTHOR INFORMATION

### Author Contributions

Conceptualization, J.M., J.G.M., I.J.M.; Data curation, J.G.M.; Formal Analysis, J.M., J.G.M.; Funding Acquisition, J.M., J.G.M., J.J.C; Investigation, J.M., J.G.M., P.S.; Methodology, J.M., J.G.M. I.J.M.; Project Administration, J.M., J.G.M., I.J.M.; Resources, J.M., J.G.M., J.J.C.; Software, J.M., J.G.M.; Supervision, J.M., J.G.M., I.J.M., J.J.C; Validation, J.M., J.G.M.; Visualization, J.M., J.G.M., P.S.; Writing – original draft, J.M., J.G.M.; Writing – reviewing & editing, J.M., J.G.M., I.J.M., J.J.C.

### Notes

J.J.C. is a consultant for Thermo Fisher Scientific and is on the scientific advisory board for Seer and 908 Devices.

## ACKNOWLEDGMENT

We thank Alex Hebert for the project idea and for helpful discussions and Evgenia Shishkova for helping PS collect the high resolution FAIMS data. This work was supported by the following NIH grants: NLM T15LM007359, R35GM142502, R35GM118110, and P41GM108538. Aspects of this research were performed using the compute resources and assistance of the UW-Madison Center for High Throughput Computing (CHTC) in the Department of Computer Sciences. The CHTC is supported by UW-Madison, the Advanced Computing Initiative, the Wisconsin Alumni Research Foundation, the Wisconsin Institutes for Discovery, and the National Science Foundation, and is an active member of the Open Science Grid, which is supported by the National Science Foundation and the U.S. Department of Energy’s Office of Science.

