## supplementary figures for "Deep Learning Predicts Non-Normal Peptide FAIMS Mobility Distributions Directly from Sequence"

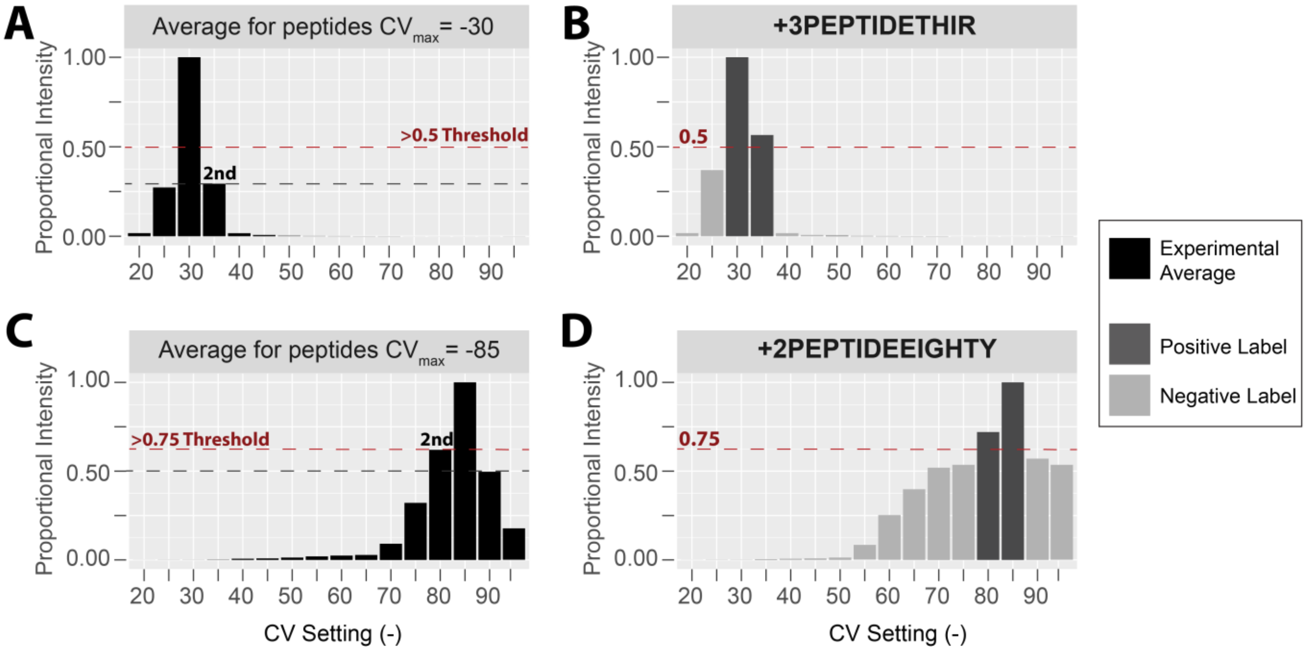


**Figure 1. True Positive CVmax-specific Labelling Scheme for the Random Forest.** Labelling thresholds are based on the average transmission distribution of peptides that share a CVmax. The threshold is set at either 0.5 or the average proportional intensity of the second-best compensating voltage, whichever is higher. (A) On average peptides with CVmax = -30 have a proportional intensity of 0.3 at -35 V, their second-best setting. (B) Because in this case of 0.3 < 0.5, our theoretical example peptide receives a positive label for each setting with > 0.5 proportional intensity. (C) In contrast, peptides with CVmax = -85 have an average proportional intensity of 0.75 at their second-best setting, -80 V. (D) 0.75 > 0.5 so our second example peptide receives a positive label for each setting with > 0.75 proportional intensity. This strategy leads to the theoretical example +2PEPTIDEEIGHTY receiving the two labels with the greatest transmission rather than five resulting from a 0.5 threshold.

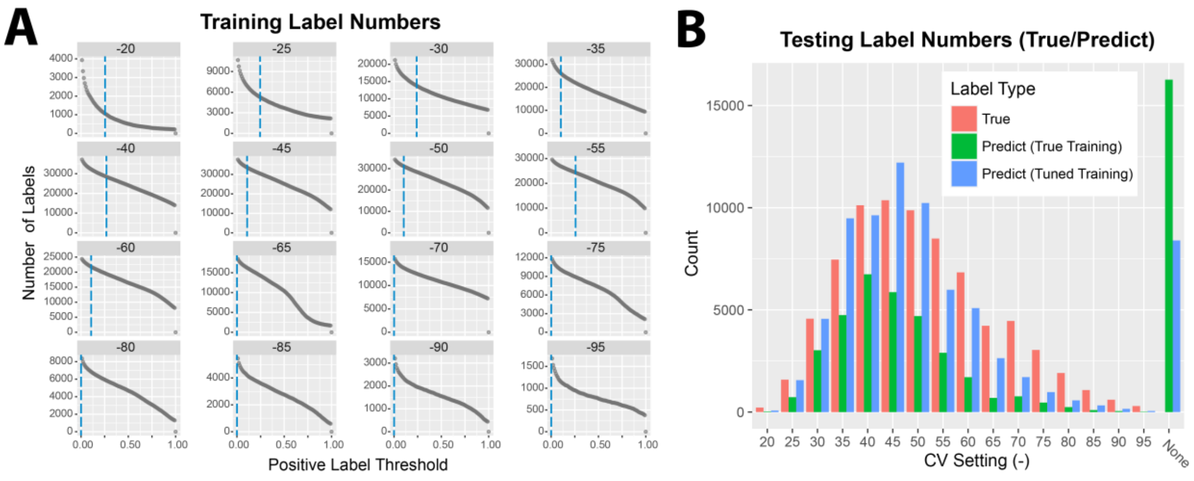


**Supplemental Figure 2. Alternate training label threshold for random forest. (A)** Number of positive labels as the labelling threshold is increased for each CV bin. Dotted green line indicates 0.5 threshold. Dotted blue line indicates lowered threshold tuned to better match prediction numbers to true label numbers in the random forest. **(B)** Number of true (red) and predicted labels that result from training the random forest using the true labelling scheme (green), or a lower, tuned labelling threshold (blue). When training with more conservative true labelling scheme, random forest substantially underpredicts labels at all bins and results in high number of peptide ions with no predicted label ("None").

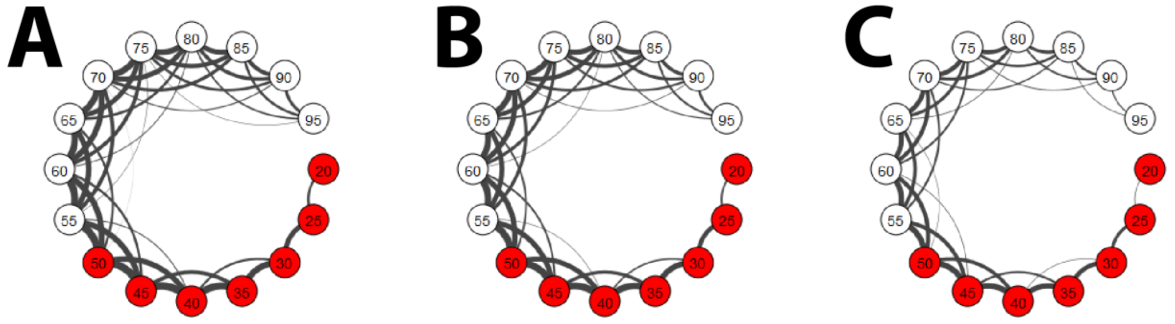


**Supplemental Figure 3. Mapping label combinations. Network plot connecting any co-occurring CV labels with thickness of connecting line indicating frequency of label combination. (A)** Combinations for all human peptides **(B)** Combinations for training human peptides **(C)** Combinations for testing human peptides. Both subsets mirror label combination distribution for human peptides overall.

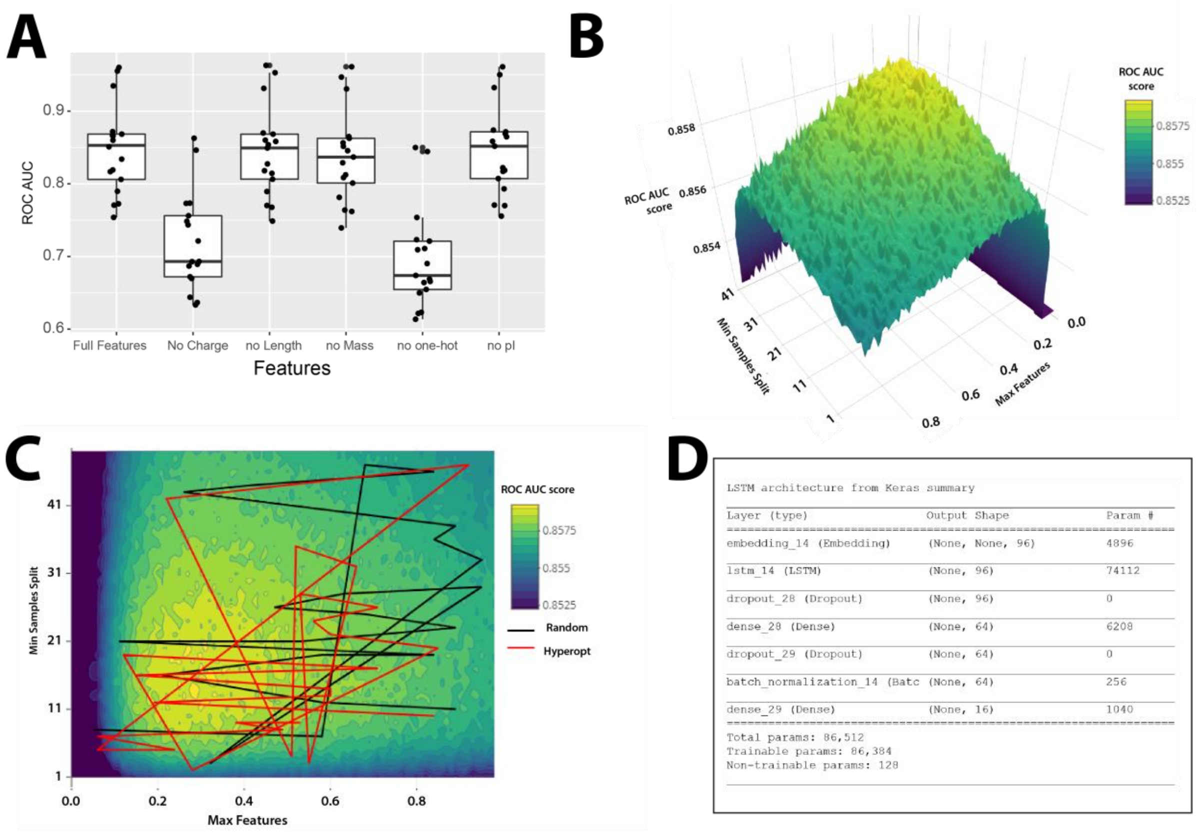


**Supplemental Figure 4. Optimization of Features and Hyperparameters. (A)** Boxplot of ROC-AUC score distribution when excluding features from the random forest model. Each point represents performance of single CV label, with bold line indicating mean score of all CVs. **(B)** Three-dimensional ROC-AUC score landscape for simple random forest model when using 100 estimators and varying two hyperparameters: min_samples_split and max_features (the two hyperparameters contributing the most to performance variability), with yellow indicating higher scores and blue indicating lower scores. Landscape contains many local maxima. **(C)** Two-dimensional ROC-AUC score landscape with a red and black line segment indicating hyperparameter space explored by the hyperopt and random search optimization functions respectively. Here we depict every other model tested by each of the two function in the interest of clarity. **(D)** Summary of the LSTM neural network architecture.

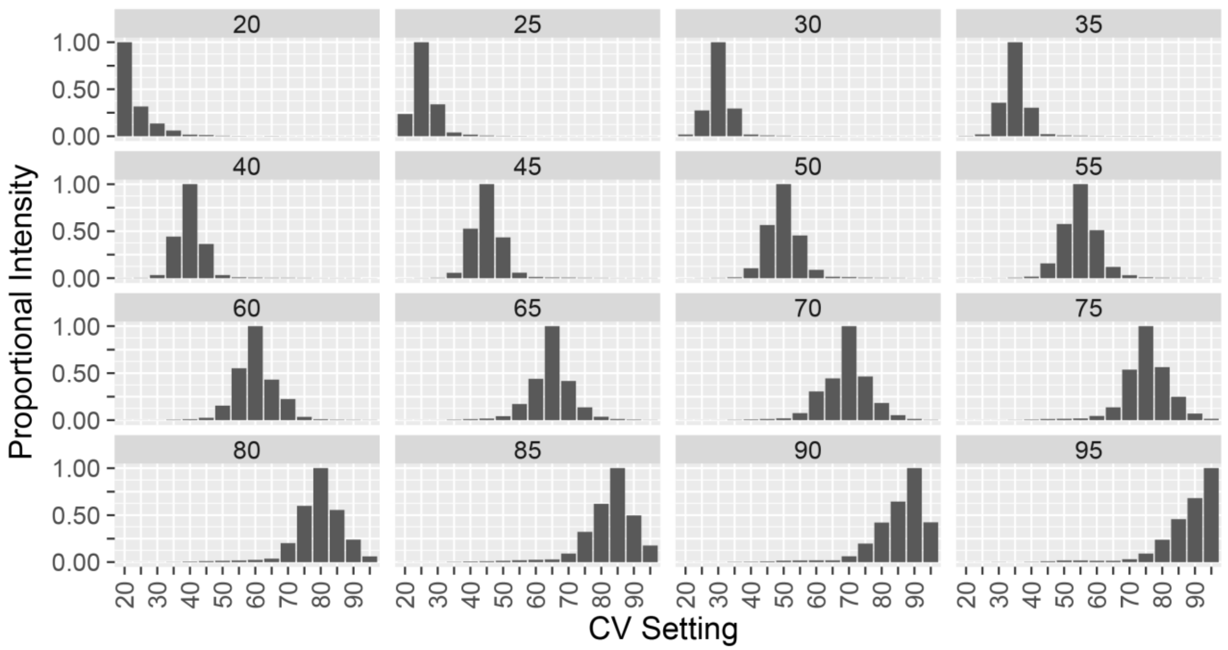


**Supplemental Figure 5. Average proportional intensity for all peptides grouped by CVmax.** Peptides with greater magnitude CVmax exhibit broader transmission distributions.

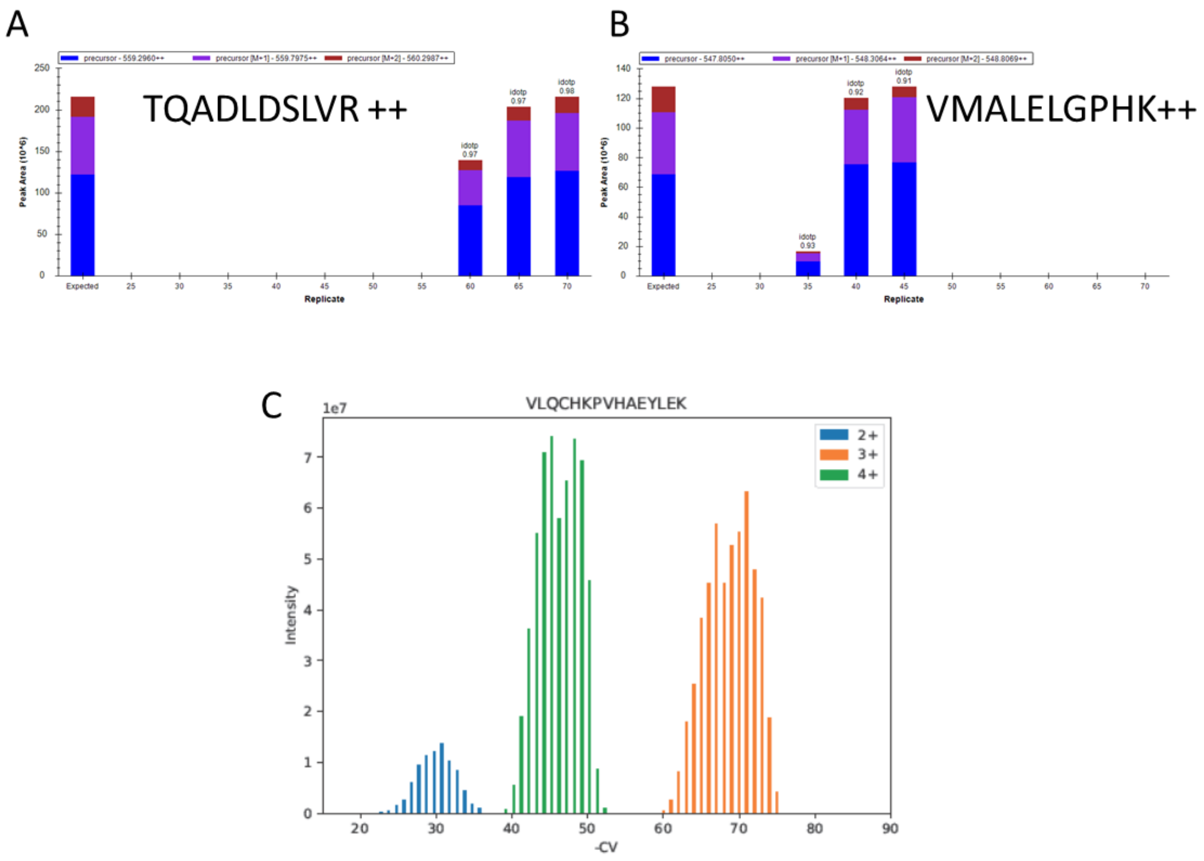


**Supplemental Figure 6: External Validation of non-normal peptide transmission distributions.** (A-B) Distributions at 5 V step size from Deng et al commonly show non-normality. (C) “High resolution” FAIMS mode plus injections at 1 V step size and 70 degrees internal electrode temperature reveal non-normal distributions of the same peptide from Figure 1C.

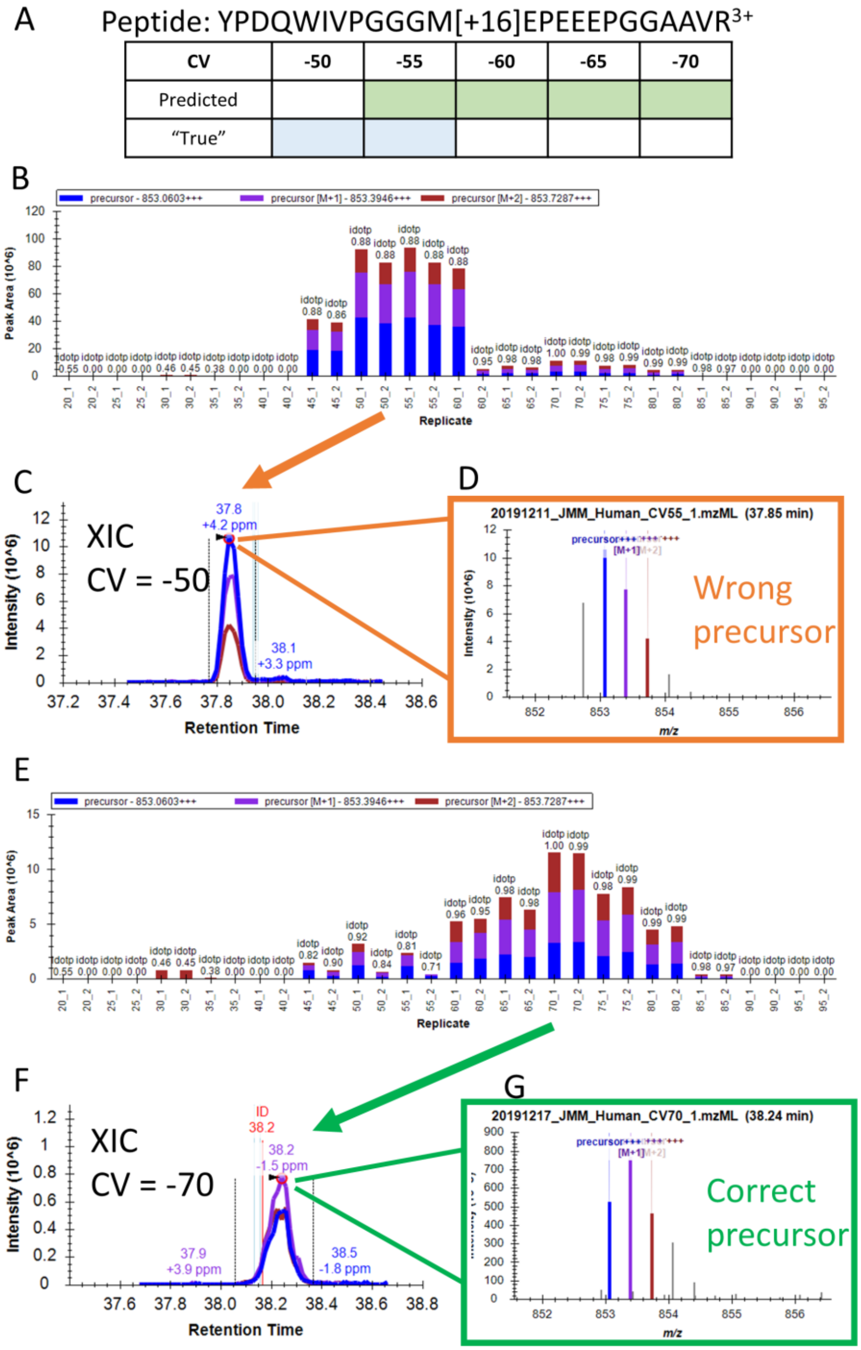


**Supplemental Figure 7: Example of "wrong" prediction due to incorrect automated peak picking in Skyline.** **(A)** Peptide sequence and table of predicted and true labels. Panels B-D show the automated peak picking by Skyline without adjustment, and E-G show after adjustment. (**B)** Peak areas for the peptide precursor across FAIMS CV duplicates. The most intense signal is at CV = -50 and CV = -55 replicates, which were assigned as true labels. The average peak areas of CV = -45 and CV = -60 duplicates would be less than ½ of the maximum and were not chosen as true labels. **(C)** Extracted ion chromatogram (XIC) of the precursor in one of the CV = -50 replicates showing good peak shape. **(D)** Peptide precursor ion envelope from the apex of the peak in **(C)** reveals that the signal comes from M+1, M+2, and M+3 isotopes of a different peptide precursor. **(E)** Peak areas over FAIMS CV duplicates after adjusting integration to the correct peak. **(F)** XIC of the peak in the CV = -70 replicate where the peptide was also identified. **(G)** Peptide precursor ion envelope at the apex of the peak in **(F)** showing the correct isotope shape.

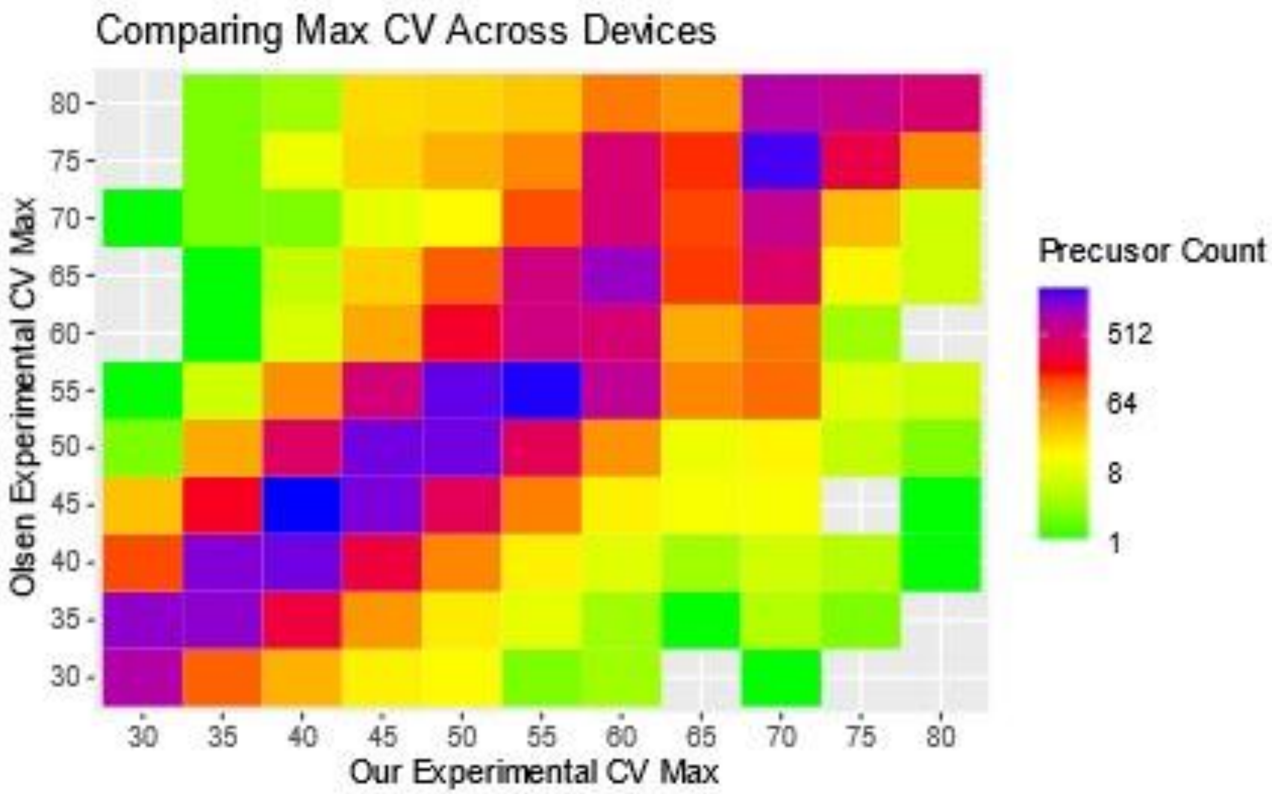


**Supplemental Figure 8:** Overlap of Max CV: Density plot indicating correlation of CVmax setting for 33,420 precursor ions commonly identified between the human dataset used here, and that from Bekker-Jensen et al.
